# Population genomics of a natural *Cannabis sativa* L. collection from Iran identifies novel genetic loci for flowering time, morphology, sex and chemotyping

**DOI:** 10.1101/2024.05.07.593022

**Authors:** Mahboubeh Mostafaei Dehnavi, Annabelle Damerum, Sadegh Taheri, Ali Ebadi, Shadab Panahi, George Hodgin, Brian Brandley, Seyed Alireza Salami, Gail Taylor

## Abstract

Future breeding and selection of *Cannabis sativa* L. for drug production and industrial purposes require a source of germplasm with wide genetic variation, such as that found in wild relatives and progenitors of highly cultivated plants. Limited directional selection and breeding have occurred in this crop, especially informed by molecular markers. Here, we investigated the population genomics of a natural cannabis collection of male and female individuals from differing climatic zones in Iran. Using Genotyping-By-Sequencing (GBS), we sequenced 228 genotypes from 35 populations. The results obtained from GBS were used to perform association analysis identifying links between genotype and important phenotypes, including inflorescence characteristics, flowering time, plant morphology, tetrahydrocannabinol (THC) content, cannabidiol (CBD) content and sex. Approximately 23,266 significant SNPs of high quality were detected to establish associations between markers and traits, and population structure showed that Iranian cannabis plants fall into five groups. A comparison of Iranian samples from this study to global data suggests that the Iranian population is distinctive and, in general, is closer to marijuana than to hemp, although some populations in this collection are closer to hemp. The GWAS results showed that novel genetic loci, not previously identified, contribute to sex, yield and chemotype traits in cannabis and are worthy of further study.

## Introduction

*Cannabis sativa* L. (cannabis) from the Cannabinaceae family can be used as a source of both pharmacologic drugs for the treatment of tumors, schizophrenia and other medical conditions, but also as fiber and oil, depending on the quantities of tetrahydrocannabinol (THC) and cannabidiol (CBD) within any particular plant, landrace or cultivar (Fischedick J, 2015; Andre et al., 2016). *C. sativa* has a long history of cultivation, and it has been suggested that the global cannabis market may be valued annually at over $300 billion in coming years, as many US states and global nations de-regulate the use of this plant-based chemical for pharmaceutical and recreational use (Weiblen et al., 2015; Grassa et al., 2021; Mostafaei Dehnavi et al., 2022). Much remains to be discovered concerning the diversity of genetic and chemical signatures across the species of *C. sativa,* and it seems likely that wild populations of previously uncharacterized *C. sativa* can provide a valuable source of natural genetic variation as foundational resources for future directed breeding programs (Zhang et al. 2020; Mostafaei Dehnavi et al. 2022). The plant is an annual species, primarily dioecious and exhibits high levels of heterozygosity. It has a diploid genome (2n=20) estimated to be 843 Mb for male plants and 818 Mb for female plants. Additionally, the species is believed to possess almost 30,000 genes (Jenkins and Orsburn, 2019b, 2019a; Henry et al., 2020; Hurgobin et al., 2021).

Depending on type and cannabinoid yield, in particular, THC: CBD ratio, this species can be defined as industrial hemp (a major source of textiles, food, and oilseed) or marijuana (medical cannabis or a recreational drug) (Piluzza et al., 2013; Ren et al., 2021; Mostafaei Dehnavi et al., 2022). There is significant potential in the development of cannabis and its derivatives, particularly CBD, in the treatment of melanoma, a type of skin cancer and epilepsy-related syndromes such as Lennox-Gastaut syndrome and Dravet syndrome (Bachari et al., 2020; Brunetti et al., 2020).

Cannabis breeding to date has been mostly outside of the public domain; therefore, the true genetic diversity of commercial varieties is unknown (Hurgobin et al., 2021). Unraveling the genetic information of natural cannabis populations facilitates breeding programs for different industrial and medical purposes (Mostafaei Dehnavi et al., 2022). Genotyping by sequencing (GBS) is a highly multiplexed and high-throughput method for determining the genetic structure of an individual or a population (Elshire et al., 2011; Sonah et al., 2013). This technique has played a significant role in the advancement of our understanding of the genetic diversity, evolution and breeding of cannabis (Sawler et al., 2015). Cannabis, as a complex plant species, has a high degree of genetic diversity, which has made it an ideal candidate for GBS studies. One of the main goals of GBS studies in cannabis is to understand links between phenotype and genotype and identify the genetic markers associated with important and complex traits (Pootakham et al., 2015).

By phenotyping a large number of plants, researchers can identify the genetic variations that are associated with specific traits and ultimately help to improve the breeding process. The phenotyping of traits such as sex, flowering time, cannabinoids production, flower structure, agronomic-related and disease and pest resistance is crucial for the cannabis industry (McKernan et al., 2020; Petit et al., 2020a; Pépin et al., 2021). Accurately identifying and characterizing these traits can inform breeding programs for the development of high-quality cannabis varieties with desirable traits. Early sex determination and the identification of molecular markers associated with sex are critical tools for cannabis growers and breeders looking to produce high-quality and high-yielding crops (Faux et al., 2016; Prentout et al., 2019). Furthermore, understanding the flowering time of a plant is a significant characteristic for not only optimizing yield and determining harvest time, but also the fiber quality and cannabinoids produced (Toth et al., 2022). Flower structure and plant height can also affect the cultivation process and crop yield and cannabinoid production, such as THC and CBD, is a key factor in the medicinal and recreational use of cannabis (Gonçalves et al., 2019; Deguchi et al., 2022; Mostafaei Dehnavi et al., 2022). Therefore, the precise phenotyping of these traits can improve crop management, increase yield and enhance the overall quality of cannabis (Woods et al., 2021).

Another important application of GBS in cannabis is in the identification and characterization of landrace cultivars. These cultivars have unique genetic profiles and are important sources of diversity for breeding programs (Mostafaei Dehnavi et al., 2022). GBS studies have helped identify the genetic relationships between landraces and characterize their genetic diversity (Soorni et al., 2017).

Genetic investigation of different natural resources assists in the development of pre-breeding and the identification of new varieties for research purposes and enables the initiation of a pipeline of novel discovery toward commercialization (Gali et al., 2019; Kovalchuk et al., 2020). In this study, we conducted genotyping of both male and female cannabis genomes to enhance our comprehension of cannabis sex evolution, as well as cannabinoid expression. We used GBS to characterize natural cannabis plant material obtained from various regions of Iran. We employed a set of 23,266 significant SNP markers, which were linked to various essential features, such as THC and CBD content (which distinguishes the drug from the hemp chemotype), sex expression, flowering time, female inflorescence features and some other morpho-physiological traits such as plant height, number of nodes, number of leaves, internode length and footstalk diameter. The investigation offers insight into population structure, genetic relationship, and genetic diversity of the cannabis species.

## MATERIALS AND METHODS

### Plant material and field experiment

A total of 35 natural cannabis populations sourced from various locations in Iran, obtained from CGRC (www.medcannabase.org), were grown in the research field of the University of Tehran. The cultivation followed a randomized complete block design with three replicates per population. The separation between each block was set at 2.5 m. Within each block, three rows were arranged for each plot (population), each extending 10 m in length, and with a row spacing of 60 cm and plant spacing of 90 cm, with a total of 10 plants planted across each row. A drip irrigation system was implemented, and plants were grown in soil amended with compost and fertilized with a balanced, water-soluble fertilizer (N, P, K) (Bhattarai and Midmore, 2014). All plants were grown under natural light conditions from April to September 2019. Throughout the growth period, daytime temperatures ranged from 31-36°C and nighttime temperatures ranged from 20-24°C, along with average daytime relative humidity fluctuated between 29% and 43%, and nighttime relative humidity ranged from 47% and 65%. To counter the impact of extreme heat and high evaporation during the leaf formation phase, the site received regular irrigation of 3-4 hours. After leaf growth, irrigation occurred three times a week, each session lasting 4-5 hours. The plants were grown to maturity, at which point they were harvested. Specific details for each population are given in **Table 1**. Collection sites and various climatic zones of the studied populations, are shown in **Figure 1**.

**TABLE 1.**
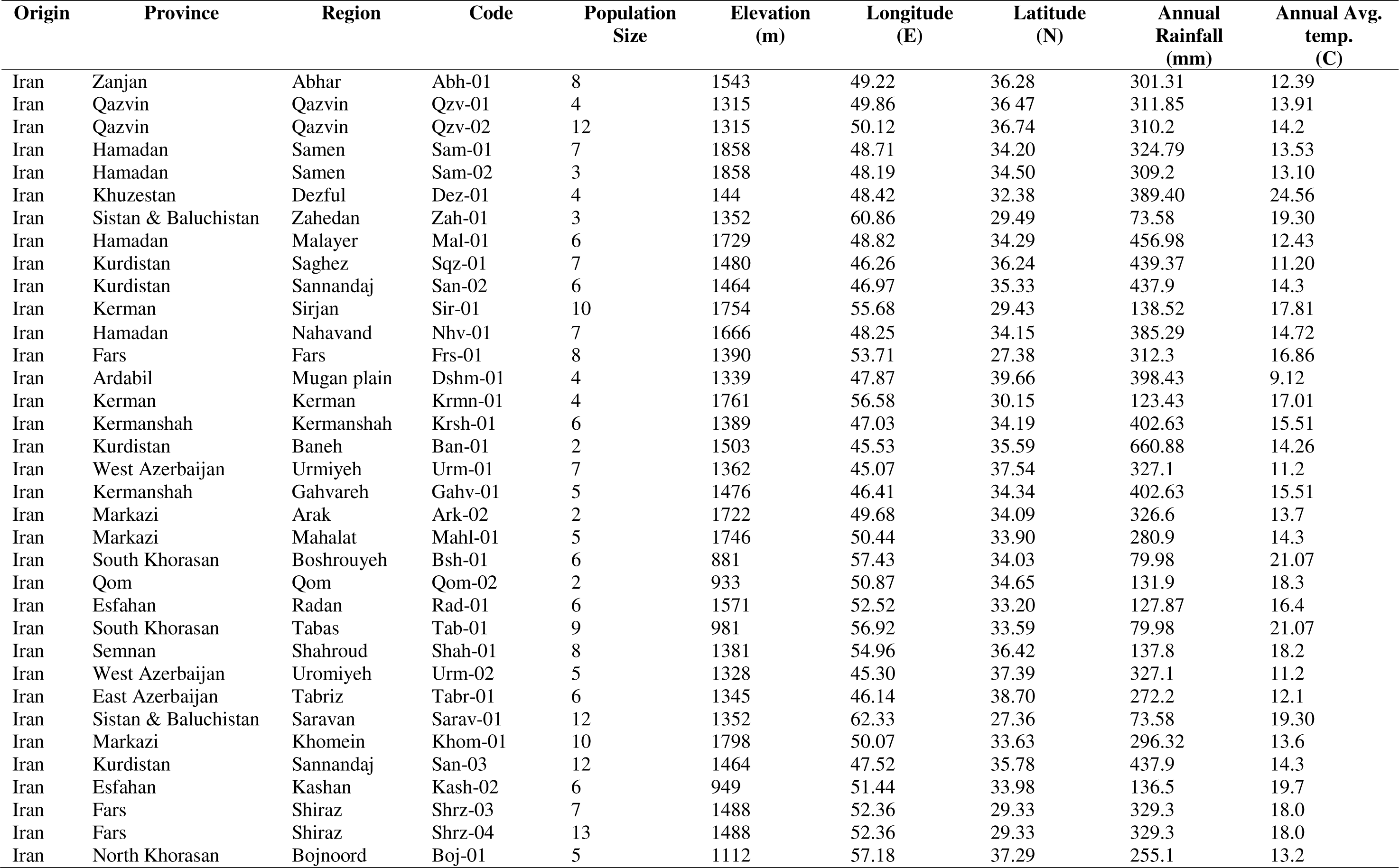
Panel of 228 *Cannabis sativa* L. and geographical and ecological parameters of the cannabis populations studied.

### Phenotyping of the GWAS Panel

For each population, randomly selected plants of the middle rows for each genotype were labeled, and the phenotypic variation of 13 traits was characterized. The number of plants employed per population varied based on availability, as highlighted in **Table 1**. Phenotypes assessed included inflorescence characteristics, flowering time, plant morphology, sex and biochemical trait analysis. Traits were (i) number of days that elapsed from germination to the initiation of flowering (DTF), (ii) the number of days from germination to appearance of approximately 50% flowering within a population (DT50), (iii) plant height (H; m) from the soil surface to the topmost terminal inflorescence before harvest, (iv) sex expression as a binary variable, i.e., male vs. female (M, F), (v) crown length, (CL; measurement of the length of the main stem from the soil surface to the lowest branch, cm), inflorescence features: (vi) inflorescence length (iL; measurement of the length of the main inflorescence in both male and female inflorescences, cm) as well as the (vii) number of lateral pistillate inflorescences (Li), (viii) internode length (intL, measurement of the length between two nodes, cm), (ix) number of nodes (nN), (x) footstalk diameter (FD) at the widest part of the base with digital calipers (cm), (xi) number of leaves (nL) and (xii) analysis of the Δ9-tetrahydrocannabinol (THC Q, %) and (xiii) cannabidiol (CBD Q, %) content of the plant material using HPLC. Air-dried pistillate inflorescences were collected prior to the seed development were analyzed for content (% dry weight (DW)) of the cannabinoid compounds Δ9-THC and CBD. Refer to our publication (Mostafaei Dehnavi et al., 2022) for information on the preparation of samples and the HPLC analysis of cannabinoids. Some characteristics, including Li, nL, DT50 and CL, were only measured in female individuals. Furthermore, it is noteworthy that all plants included in this study exhibited dioecious characteristics, with either male or female flowers. As a result, we recorded the sex expression (male or female) of each individual in every population.

### DNA Extraction, Library Preparation and Genotyping of the GWAS Panel

The GBS method was utilized to genotype the GWAS panel, following the protocol delineated by Elshire et al. (2011). At the juvenile stage before sexual differentiation, leaf tissue was collected from a labeled single plant of each population, and small segments of the tissue were placed into 2 ml vials and freeze-dried. High molecular weight DNA was isolated from approximately 25 mg of freeze-dried tissue following a modified cetyl trimethyl ammonium bromide (CTAB) protocol (Doyle & Doyle, 1987; Cullings, 1992), which included a step for RNase treatment, to remove any potential RNA contamination, as RNA can inhibit the DNA sequencing library preparation (https://dnatech.genomecenter.ucdavis.edu/faqs/which-dna-isolation-protocols-do-you-recommend-for-illumina-sequencing/). DNA extracts were quantified using a Qubit^TM^ Fluorometer (ThermoFisher Scientific). Individual DNA samples were diluted to 10 ng/μl using 0.5 M Tris-EDTA (TE) buffer, pH 8.0. As the sex of the plants was unknown, to cover all allelic variation within populations and sexes, the genomic DNA was extracted from all available plants per population. 100 ng of each genomic DNA template (in a 10 μl volume) was used for library construction using a single digestion with restriction enzyme *ApekI* and ligated to unique 4-8 sequence barcode adapters. Five μl aliquots of adapter-ligated DNA samples were pooled in a single tube to produce 96-plex libraries. The pooled DNA was PCR-amplified using *Phusion® High-Fidelity PCR Kit* (NEB*®*), followed by purification with a *Monarch®* PCR & DNA Cleanup Kit (NEB*®*). Standard experimental conditions, as described by Elshire et al. (2011), were followed for restriction, ligation, and PCR amplification. The purified DNA library was quantified and validated using a Bioanalyzer (Agilent Technologies), and the 96-plex libraries were sequenced on a single lane of Illumina HiSeq™ 4000 platform as single-end 100 (SR100) base pair reads.

### Demultiplexing, Data quality control, and Read filtering

The Stacks pipeline was used for GBS data analysis (Catchen et al., 2013). Demultiplexing and trimming the sequence reads were performed using *process_radtags* script, which trims adapter sequences and filters low-quality reads <50 bases. Samples with <100,000 reads were removed before analysis. To elucidate the genetic relationship among Iranian cannabis and marijuana and fibre type accessions, we integrated our data with two public datasets. This included marijuana data consisting of 81 samples and hemp data consisting of 43 samples originally prepared by Sawler et al. (2015) and obtained from the NCBI SRA BioProject: PRJNA285813. Additionally, we incorporated 95 cannabis samples including 70 from Iran, 2 from Afghanistan and 26 accessions provided by CGN and IPK, as previously reported by Soorni et al. (2017), and accessed from the BioProject: PRJNA419020.

### Mapping, SNP Variant Calling and SNP Filtering

We used a reference-based pipeline for sample alignment to generate consensus sequences. Trimmed sequence reads were aligned to the reference *C. sativa* ‘CBDRx’ assembly (cs10 v.1.0) as the most complete and contiguous chromosome-level assembly available at the time of analysis (Hurgobin et al., 2021; Ren et al., 2021), using the sequence alignment tool bowtie2 (http://bowtie-bio.sourceforge.net/bowtie2/manual.shtml#the-bowtie2-aligner) with the very-sensitive-local settings. This resulted in an average mapping rate of approximately 80%. The *ref_map.pl* script within the Stacks environment was utilized to call genetic variants (SNPs), and the "*populations"* program of the Stacks pipeline was used to filter the identified SNPs and estimate population genetics statistics like the fixation index (F_ST_) for genetic relationship analysis and Hardy-Weinberg equilibrium (hwe). The filtration criteria applied were as follows: requiring a locus to be present in a minimum of 10 populations for processing; setting a minimum of 50% individuals per population to process a locus for that population; necessitating a minimum of 50% individuals across populations for locus processing; and specifying a minimum minor allele frequency (MAF) of 0.05 for processing nucleotide sites at a locus. PLINK V 1.9 (Purcell et al., 2007) was used for further filtering for both datasets (derived from this study and the publicly available data). Individuals with a genotyping call rate < 99%, SNPs with a genotyping call rate < 99%, and those exhibiting significant deviations from Hardy-Weinberg equilibrium (P-value <10^-6^) were excluded from the analysis (Taheri et al., 2023). Following this filtering process, the genetic data from both groups were combined, and only the SNPs that were common to both groups were selected. The set of obtained SNPs was used for subsequent analysis, including population structure analysis, heterozygosity and F_ST_ analysis and association analysis.

### Population Structure Analysis

*Admixture* 1.3.0 (Alexander et al., 2009) was utilized to estimate the most likely number of clusters (K) into which the accessions could be grouped and their degree of admixture. The value of K that best fits the data was determined based on the lowest cross-validation (CV) error. Accessions were assigned to clusters based on the probabilities of belonging to one of the clusters derived from the matrix of contributions, Q. *Admixture* was run for each possible group number (K = 1 to 10). In addition, to visualize the genetic relationship and similarity among samples, a principal component analysis was carried out on a combined dataset of 431 samples.

The analysis utilized ggplot2 (V3.4.4) for plotting (Villanueva and Chen, 2019), plotrix (V3.8.4) for zooming the plot (Lemon et al., 2015), and tidyverse (V2.0.0) for eliminating duplicate samples (Wickham et al., 2019) in R (V4.3.1). Following the quality control filtration process, this dataset consisted of 196 Iranian samples from the current study, 93 cannabis samples previously studied by Soorni et al. (2017), as well as 47 individuals from a hemp population and 95 individuals from a marijuana population studied by Sawler et al. (2015).

### Heterozygosity and F_ST_ analysis

Heterozygosity was estimated for each individual using PLINK V1.90 and then averaged within each group (Purcell et al., 2007). Additionally, we used R (V4.3.1; R Core Team, 2018) to generate plots.

Due to the limited number of individuals within some populations, we classified the studied Iranian populations into four larger groups based on their geographical distribution and associated climatic patterns. These groups include east and southeast, northeast, south and west and northwest populations. The F_ST_ value was calculated by fsthet package (Flanagan and Jones, 2017) in R (V4.3.1) for each pair of populations to measure genetic differentiation among populations. F_ST_ analysis was conducted three times: firstly, among the four geographic populations of Iranian samples; secondly, using combined data from this study and two public datasets containing previously studied Iranian samples, as well as other marijuana and hemp populations (NCBI SRA BioProject: PRJNA419020 and PRJNA285813), and in the third analysis, all Iranian populations from this study were treated as a single population and combined with two above-named public collections. The F_ST_ plots were created in R (V4.3.1) using the qqman package (Turner, 2014) to visualize the relationships between populations based on the F_ST_ values. We then conducted a thorough analysis of significant SNPs for each pair to identify the specific SNP markers contributing to the observed differences.

### IBD Test

We employed PLINK V1.90 software to conduct pairwise IBD analysis, investigating first-degree and second-degree relationships among individuals by assessing the proportion of SNPs where zero, one, or two shared IBD alleles were present, represented by Z0, Z1, and Z2, respectively. Subsequently, relatedness was quantified using the PI_HAT parameter, indicating the proportion of SNPs in IBD between individual pairs (Arab et al., 2019).

### Association Analysis

To identify the associations between genetic varaints and trait performance, GWAS was carried out to estimate SNP effects. Studies indicates that employing a linear mixed model that incorporates population and family structures is currently the most effective approach for mitigating the impact of population stratification (Price et al., 2010). The statistical model used for GWAS analysis is based on the mixed linear model as follows:

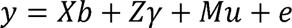

Where y is a vector representing the phenotype, b is a vector representing fixed effects (group), γ is a vector representing fixed effects of markers, and u is a vector representing random effects (e.g., PC1, PC2, and PI_HAT). X, Z, and M are matrices relating observations to the effects of fixed factors, fixed SNP effects, and random genetic effects, respectively, and e is a vector representing random residuals with e ∼ N(0, I σe2).

The analysis initially utilized a mixed linear model through GCTA V1.94.1 software (Yang et al., 2011). Additionally, for undertaking GWAS for sex as a qualititative trait, we employed PLINK V1.90 with case and control anaylsis. Subsequently, we applied Bonferroni testing using PLINK’s --assoc, --perm, and --adjust functions. Finally, Manhattan plots and the Q-Q plots were constructed in R (V4.3.1) using the qqman package (Turner, 2014) to visualize the genome-wide association signals.

Furthermore, GCTA V1.94.1 software was utilized to measure the SNP-based heritibility (h^2^) for each trait (Zhu and Zhou, 2020). The variance of total additive genetic effects is defined as 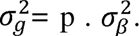 The GCTA software was used to estimate the variance components 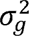 and 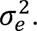 The SNP heritability is estimated as follows:

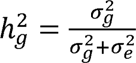

### Functional annotation of candidate SNPs

In addition, functional annotation of the candidate SNPs identified in the GWAS analysis was performed. For each trait, the significant markers were compared and annotated using the annotated reference genome (cs10, GCF_900626175.1, https://www.ncbi.nlm.nih.gov/genome/annotation_euk/Cannabis_sativa/100/) and the annotated genes were identified using the NCBI Genome Data Viewer (https://www.ncbi.nlm.nih.gov/genome/gdv/browser/genome/?id=GCF_900626175.2) (Jenkins and Orsburn, 2019b).

## RESULTS

### Phenotyping, genotyping and data quality control

A normal distribution was observed for all traits except for DT50, THC Q, CBD Q, nL, and DTF (**Figure 2**). Sex was not included as it is not feasible due to its non-quantitative nature. We observed great phenotypic variation within our cannabis populations. The footstalk diameter exhibited substantial diversity, ranging from 1.62 to 3.66, with an average of 2.54, showing the highest coefficient of variation at 31.32%. In contrast, THC and CBD concentrations displayed narrower variations, ranging from 1.17 to 3.13% for THC and 0.96 to 6.69% for CBD, with averages of 1.96% and 1.65%, respectively. Consequently, they exhibited the lowest coefficients of variation at 0.51% and 0.52%, respectively (**Table 2**). The phenotypic data collected for the GWAS panel, along with the SNP-based heritability (h^2^) measurments are summarized in **Table 2**. A total of 1.456 billion reads of 100 bp length were obtained from the four sequencing lanes on HiSeq^TM^ 4000. After quality control and trimming for barcode adapter sequences, an average of approximately 4 million reads per sample were retained for the first (3,883,851 reads/sample; 80.6% of original reads), second (3,854,236 reads/sample; 82.8%), and third (3,935,551 reads/sample; 79.5%) libraries, while approximately 10 million reads (10,774,545 reads/sample; 64.3%) remained for the fourth library, which included only 35 pooled samples. The number of reads retained per sample ranged between 108K and 104M reads, with samples with <100K reads removed. The average quality scores, Q30 ratio and guanine–cytosine (GC) content of the reads were ∼39, ∼96% and 44.1%, respectively. On average, 80.11% of the reads aligned with the *C. sativa* cs10 reference genome (Hurgobin et al., 2021). Differences in sequencing depth across regions, excessive PCR amplification, short read length, or issues with the sequencing platform may have contributed to the variations observed in the percentage of reads mapped to the reference. Specifically, the minimum mapping percentage recorded was 50.09%, which was observed in the case of sample ID 218, representing one of the individuals from population Bsh-01. Following filtration, a total of 23,266 high-quality SNPs were identified across Iranian samples, which were subsequently selected for the analysis of population structure and marker-trait association. Additionally, a set of 25,112 informative SNPs were retained after applying filtration to the combined datasets, which included the previously sequenced Iranian samples and collections of hemp and marijuana.

**TABLE 2.**
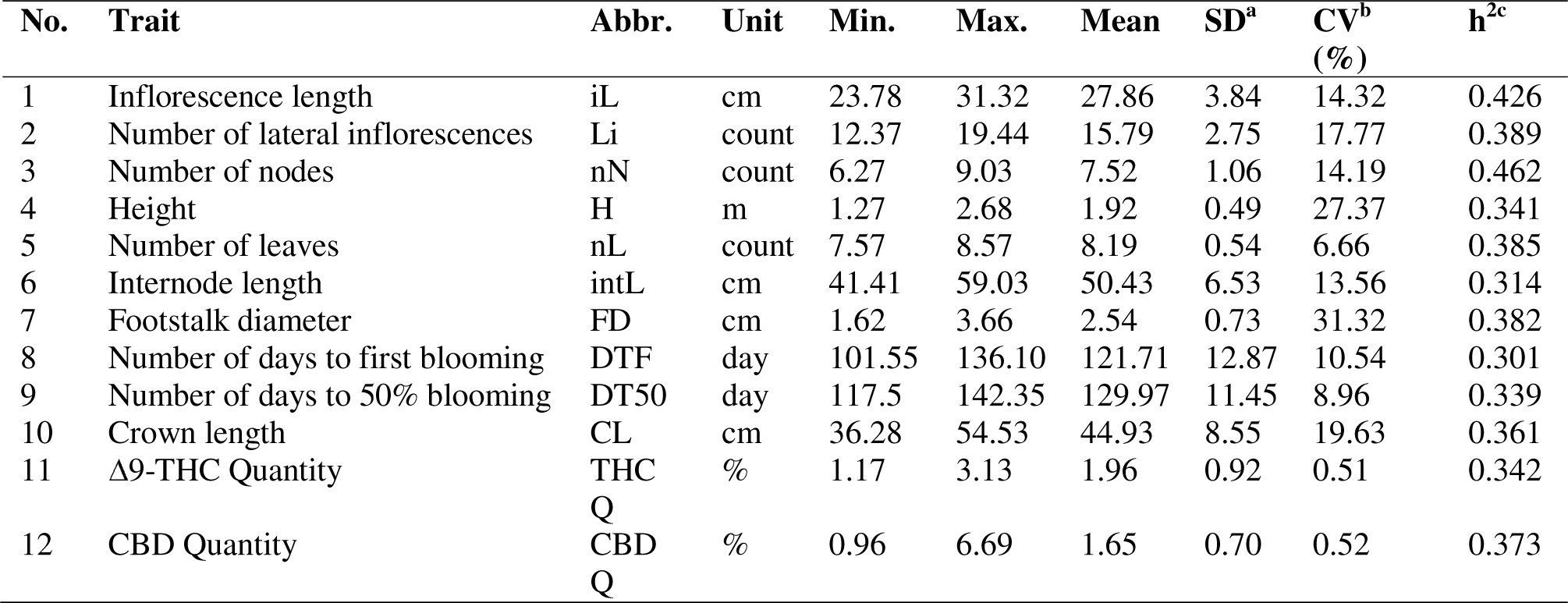
Characteristics studied in the cannabis populations (sex trait has not been reported here as it is not a quantitative trait). ^a^SD -(standard deviation), ^b^CV - (coefficient of variation), estimated as the ratio of the standard deviation to the mean of all populations and ^c^h^2^- (SNP-based heritability), the proportion of phenotypic variance explained by all measured SNPs.

### Population Structure Analysis

Principal Component Analysis (PCA) findings indicated a high level of genetic similarity among these populations (Figure 3A). The Sqz-01 population and some individuals of San-02 failed to group with other clusters (Figure 3A). Admixture’s cross-validation procedure was used to determine the most likely number of genetic groups (K). The population structure of studied samples was described by testing the probable number of clusters (K) from 1 to 10, with K= 5 selected as the optimal representation of ancestral populations based on the lowest cross-entropy criterion and visualized using a Q estimates bar plot (Figure 3B-C). The PCA analysis of integrated data with two public datasets revealed that while the Iranian samples exhibited distinct genetic differences from the hemp and marijuana populations, they showed generally closer genetic proximity to the marijuana population. However, some individual Iranian samples exhibited a closer genetic resemblance to hemp (Figure 4A). These findings were further supported by the dendrogram plot, which indicated that the genetic distance between the Iranian samples from this study and the previously studied marijuana population was smaller than the distance between the Iranian samples from the current study and the previously studied hemp population (Figure 4B).

### Genetic variation and differentiation

To assess the genetic differentiation among Iranian cannabis populations, we calculated the genetic differentiation parameter (F_ST_) as well as observed and expected heterozygosity for each pairwise comparison. Initially, we generated six F_ST_ plots to compare distinct populations of Iranian samples (**Supplementary Figure 1**). The observed heterozygosity (H_o_) for these groups was found as 0.25 (for the east and southeast population), 0.204 (for the northeast population), 0.214 (for the south population), and 0.228 (for the west and northwest). Corresponding, the respective expected heterozygosity (H_e_) values were obtained as 0.288, 0.264, 0.286, and 0.296, while the estimated minor allele frequency (MAF) were 0.204, 0.19, 0.203, and 0.208. The details of observed heterozygosity, expected heterozygosity, and minor allele frequency are provided in **Figure 5**.

Our study exhibited higher heterozygosity than that earlier study on Iranian samples observed by Soorni et al. (2017). Our results showed a similar level of heterozygosity to the hemp accessions studied by Sawler et al. (2015), while the earlier Iranian sample study reported the heterozygosity, which is more similar to marijuana accessions studied by Sawler et al. (2015).

**Supplementary Figures 1**, **2**, **3** present F_ST_ Manhattan plots and correlation plots for all possible pairings of these populations. Significant SNP markers along with their corresponding loci, gene annotations, and the F_ST_ values across all F_ST_ analyses are provided in **Supplementary Tables 2-18**.

The pairwise comparisons among the geography-based Iranian populations in this study revealed that the east and southeast population and northeast populations had the highest number of SNP markers (95 SNPs) (**Supplementary Figure 1C**; **Supplementary Table 2**), followed by the northeast population and south population with 89 SNP markers which related to 32 specific loci (**Supplementary Figure 1F**; **Supplementary Table 3**). On the other hand, the east and southeast population and southern population exhibited the lowest number of SNPs (29 SNPs) that were linked to five specific loci (**Supplementary Figure 1D**; **Supplementary Table 5**).

When comparing Iranian populations with global collections, in contrast to the hemp collection, the marijuana collection exhibited a larger number of SNP markers concerning the geography-based Iranian populations. Among these comparisons, the northeast population of Iran demonstrated the highest number of SNP marker (134 SNPs) spanning 36 loci in comparison to the marijuana collection (**Supplementary Figure 2A**; **Supplementary Table 13**). Furthermore, the northeast population emerged with the highest number of SNP markers (112 SNPs) linked to 49 loci when compared to the hemp collection (**Supplementary Figure 2C**; **Supplementary Table 9**). Overall, the northeast population of Iran appeared to be more distinct not only from other populations within Iran but also from both the hemp and marijuana collections.

In a separate analysis, pairwise comparisons were conducted between the Iranian samples from this study as a single population and previously studied Iranian samples, as well as hemp and marijuana collections. It was found that there were less SNP markers shared between Iranian samples and hemp (107 SNPs) (**Supplementary Figure 3A**; **Supplementary Table 16**) than between Iranian samples and the marijuana collection (113 SNPs) (**Supplementary Figure 3B**; **Supplementary Table 17**). In both of these comparisons, a total of 37 loci were found to exhibit differences (**Supplementary Figure 3A, B**; **Supplementary Tables 16, 17**). Of these, six loci were identified as being common to both comparisons, including LOC115694687, which encodes separase (RefSeq accession: XM_030621771.1), LOC115707169 encoding for histone deacetylase 2 (RefSeq accession: XM_030635038.1), LOC115707184 encoding for spidroin-2 (RefSeq accession: XM_030635066.1), LOC115707237 encoding for serine/threonine protein phosphatase 2A 55 kDa regulatory subunit B (RefSeq accession: XM_030635130.1), LOC115725648 with an uncharacterized description (RefSeq accession: XM_030655228.1), and LOC115725736 encoding for partner of Y14 and mago (RefSeq accession: XM_030655329.1).

It is noteworthy that the comparisons unveiled 134 SNP markers associated with 51 loci that exhibited differences between the Iranian samples of this study and the previously studied Iranian samples. These SNPs were distributed across chromosomes 1 (n=5), 2 (n=11), 4 (n=31), 5 (n=27), 7 (n=6) and 10 (n=54) (**Supplementary Figure 3C**; **Supplementary Table 18**). This suggests the existence of genomic differences between these two sample sets.

In the first F_ST_ analysis conducted among the four geographical-based Iranian populations, the highest and lowest F_ST_ values were observed for the east and southeast: northeast pair (F_ST_= 0.09) and northeast: west and northwest pair (F_ST_= 0.024), respectively. A higher F_ST_ value indicates greater genetic differences between populations. Moving on to the second F_ST_ analysis, which incorporated global data, the pairs of marijuana: northeast demonstrated the lowest F_ST_ value (F_ST_= 0.06), while the pair of hemp: east and southeast exhibited the highest F_ST_ value (F_ST_= 0.17). These results suggest that Iranian populations displayed higher F_ST_ values compared to hemp rather than marijuana. Furthermore, in the last set of F_ST_ comparisons, it was found that the F_ST_ estimation between Iranian samples as a single population and the hemp collection (F_ST_= 0.086) was higher compared to the F_ST_ value between Iranian samples and the marijuana population (F_ST_= 0.062). Additionally, the F_ST_ value between the Iranian samples from this study and the Iranian samples from previous studies was 0.015. Overall, these findings indicate that Iranian populations are genetically closer to marijuana than hemp. Moreover, the SNP markers that revealed differences within Iranian populations were predominantly concentrated on Chromosomes 3, 4, and 1, indicating potential genomic regions that contribute to genetic variation and population differentiation. These specific chromosomes may harbor genes or regulatory elements that play a significant role in shaping the unique genetic landscape of Iranian populations, warranting further investigation to uncover the underlying genetic mechanisms and potential functional implications of these variations.

### Genotype and phenotype associations

A total of 191 significant SNP associations were detected for the investigated traits. Among these, 47 SNPs were identified within annotated genes. Overall, the study revealed a total of 47 candidate loci related to the traits under investigation, out of which seven genes remain uncharecterized. Detailed information for all identified significant SNPs, is provided in **Supplementary Table 1** and **Table 3** with associated candidate loci, and annotation information presented specifically for those SNPs located within annotated genes. Specifically, 18 SNPs were found to be associated with H (**Figure 6A**). Among these SNP markers, five markers, two on chromosome 3 (chr3_281101_71, chr3_288389_58), two on chromosome 4 (chr4_327964_19, chr4_382581_26), and one on chromosome 2 (chr2_84187_7) were linked to annotated genes (**Supplementary Table 1**). 19 SNP markers were found to be associated with the nN, while 17 SNP markers were associated with the Li. Specifically, for the nN, there were five SNP markers located on chromosomes 2, 4, 5 and 9 that were located within annotated genes (**Figure 6B**). (**Supplementary Table 1**). Similarly, for the Li, there were two markers on chromosomes 5 and 9 that were identified within annotated genes (**Figure 6C**; **Supplementary Table 1**). Among 10 markers identified as being associated with the iL, a total of four SNP markers positioned on chromosomes 2, 9 and 10 were found to be related to genes (**Figure 6D**; **Supplementary Table 1**).

**TABLE 3.** Functional annotations of the significantly associated SNPs for sex trait in Iranian cannabis collection

| CHR | SNP ID | Position | P-value | MAF | Gene ID | RefSeq Accession | Gene Annotation |
| --- | --- | --- | --- | --- | --- | --- | --- |
| 1 | 786516_36 | 4175218 | 0.000969 | 0.33 |  |  |  |
| 1 | 786516_49 | 4175231 | 0.000969 | 0.33 |  |  |  |
| 1 | 786516_53 | 4175235 | 0.000969 | 0.33 |  |  |  |
| 2 | 78363_79 | 87241293 | 0.000352 | 0.42 | LOC115721262 | XM_030650525.2 | uncharacterized LOC115721262 |
| 2 | 109511_15 | 101011485 | 0.000414 | 0.12 |  |  |  |
| 9 | 585232_75 | 31767032 | 0.00059 | 0.24 | LOC115722331 | XM_030651522.2 | CRM-domain containing factor CFM3A, chloroplastic/mitochondrial |
| 10 | 666147_74 | 42986208 | 0.000558 | 0.27 |  |  |  |
| 10 | 689517_88 | 55869380 | 0.000621 | 0.33 |  |  |  |

CHR, chromosome; MAF, minor allele frequency.

Significant associations were observed between flowering time-related traits, including DTF and DT50 (**Figures 6E**, **6F**). 16 markers were associated with DTF, while eight markers were found to be associated with DT50 (**Figures 6E and 6F**; **Supplementary Table 1**). The SNPs associated with DTF were found to be linked with candidate loci, including glutamate--glyoxylate aminotransferase 2, YTH domain-containing protein ECT4, ribonuclease H2 subunit B, and one uncharacterized gene. As for the DT50 trait, only one marker located on chromosome 5, encoding the putative pentatricopeptide repeat-containing protein At2g02150, was identified (**Supplementary Table 1**).

Among 17 SNP markers that were detected to be associated with the FD, six SNPs, two on chromosome 5 and one each on chromosomes 2, 3, 4, and 10 were linked to annotated genes (**Figure 6G**; **Supplementary Table 1**). Also, four SNP markers on chromosomes 3 (n=2), 5 (n=1) and 6 (n=1) from a total of seven markers were detected to be associated with annotated genes for the nL trait (**Figure 6H**; **Supplementary Table 1**).

For the intL trait, seven markers positioned on chromosomes 3, 4 and 5 were associated with genes (**Figure 7A**; **Supplementary Table 1**). These markers were part of a total of 35 markers identified to be linked with this trait. Regarding the CL trait, among 12 markers that were found to be associated with it, five markers-three on chromosome 3 and two on chromosome 4 were related to annotated genes (**Figure 7B**; **Supplementary Table 1**).

Eight markers, two on chromosome 1, two on chromosome 3, two on chromosome 8 and one each on chromosomes 7 and 10 were detected to be associated with THC Q trait (**Figure 7C**; **Supplementary Table 1**). Furthermore, in the case of CBD Q, a total of 24 SNPs were identified. These markers were distributed across whole genome (**Figure 7D**). The candidate loci identified to be associated with CBD Q trait included the berberine bridge enzyme-like D-2 (chromosome 7), 3-ketoacyl-CoA synthase 19 (chromosome 1), serine/threonine-protein kinase AtPK1/AtPK6 both (chromosome 3), fimbrin-2 (chromosome 4), endonuclease MutS2 (chromosome 5), small nucleolar RNA R71 (chromosome 2), transcription factor MYB14 (chromosome 9), and one uncharacterized locus (chromosome 4) (**Supplementary Table 1**).

It appears that THC and CBD concentrations have complex genetic architectures that extend beyond the already identified cannabinoid synthase genes and are distributed across various chromosomes of the whole genome.

The SNPs associated with sex determination were primarily distributed on the sex chromosome (cs10 v.1.0 chromosome 1 and cs10 v.2.0 chromosome 10), previously identified by Prentout et al. 2019 and McKernan et al. 2020, but additional SNPs were also identified at different positions. Eight SNP markers discovered for this trait, which are located on chromosomes 1 (n=3), 2 (n=2), 9 (n=1) and 10 (n=2). Among these, two SNPs located across chromosomes 2 and 9 were found to be specifically linked to candidate genes associated with sex (**Table 3**; **Figure 7E**). The Manhattan plots, along with the corresponding quantile-quantile (Q-Q) plots for each trait, were presented in Figures **6A-6H** and **7A-7E**.

## DISCUSSION

This study adds significantly to our limited understanding of the population genomics of *Cannabis sativa* and provides novel insights into gene-trait associations for a natural population originating from contrasting climatic zones across Iran. The scarcity of such studies likely results from the historical constraints in accessing wide cannabis populations that capture natural genetic diversity (Campbell et al., 2019; Petit et al., 2020b; Hurgobin et al., 2021; Mostafaei Dehnavi et al., 2022). These insights are new, with few previously published GWAS and population genomics studies available for this largely undomesticated crop, and they add important knowledge to our developing understanding of the genomics of medicinal and industrial characterization of the crop. They will help underpin directed breeding programs to enhance traits of interest for commercial production (Hurgobin et al., 2021) and also add new insight to the population genomics and domestication history recently reported for *C. sativa* (Ren et al., 2021) that did not include data from the wild populations of Iran, such as those reported and characterized here.

The investigation of genetic variation between domesticated and natural populations is crucial for comprehending the patterns of local adaptation and identifying the genetic sources of desirable characteristics (Mostafaei Dehnavi et al., 2022). Studies on wild landraces are emerging, for example, for the wild population of Cannabis in China (Chen et al., 2022), which revealed five distinct groups and the population genomics for this important site of Cannabis domestication is important, since China is believed may possibly be one of the main centers of origin for this crop (Zhang et al., 2018; Gao et al., 2020).

Genotyping of this Iranian wild collection using GBS allowed us to conduct genetic diversity (genetic distance), population structure and genome-wide association analysis among these native populations, such as has been performed in wild (feral) cannabis collections as well as in other species, where wild progenitor populations have been used to inform breeding (Piluzza et al., 2013; Labate et al., 2014; Arbelaez et al., 2015; Ivanizs et al., 2019; Schwabe et al., 2021; Busta et al., 2022; Muli et al., 2022; Blois et al., 2023; Pandey et al., 2023). The results of population structure revealed the presence of five clusters, in contrast to the two genetic clusters reported in the previous investigation of Iranian cannabis populations conducted by Soorni *et al*. (2017). The estimation of K is influenced by two crucial factors: the number of populations and the genetic dispersion among them (Zhang et al., 2022). The observed variation may be attributed to the inclusion of a broader range of populations situated in diverse climatic zones and a larger sample size per population. Additionally, the Sqz-01 population did not form a cluster with other population groups. As previously stated, this population stands out due to distinctive morphophysiological characteristics, such as dwarf stature, early flowering, and compact inflorescence, which differentiate it from the other populations (Mostafaei Dehnavi et al., 2022).

Due to its predominantly wind-pollinated dioecious nature, cannabis is a highly heterozygous and outcrossing species (Lynch et al. 2016; Welling et al. 2020). Sawler *et al*. (2015) noted greater heterozygosity in hemp than in marijuana. Heterozygosity across distinct geographically-based Iranian populations showed a similarity in the heterozygosity to that of hemp accessions studied by Sawler et al. (2015), while the earlier study on Iranian samples reported an average heterozygosity which is more similar to that observed in marijuana. This is despite the fact that Lynch *et al*. (2016) observed a significant rise in heterozygosity within drug-type varieties compared to hemp varieties. These distinctions highlight the complex interplay between genetic backgrounds and environmental factors, resulting in diverse heterozygosity patterns within cannabis populations (Zhang et al., 2018). The expected heterozygosity values were higher than the observed heterozygosity in all four geography-based groups of Iranian populations, possibly indicating the impact of inbreeding and reduced genetic variability (Radosavljević et al., 2015). However, the limited sample size of our cannabis collection may also have contributed to these results and increasing the sample size of the population could further improve the results (Nei, 1978; Stevens et al., 2007; Arab et al., 2019; Schmidt et al., 2021).

Lower F_ST_ values between Iranian population pairs ranging from 0.02 to 0.06 indicate no strong genetic differentiation among these populations (Wrights, 1978). This phenomenon could be attributed to the distribution of pollen and seeds and the gene flow between these areas (Cheng *et al*., 2020). These seed exchanges could also have been facilitated by human activity, particularly for distant locations, as well as by wind pollination and seed dispersal by bird movements (Kitada et al., 2021). In the earlier investigation by Soorni *et al*. (2017), it was similarly noted that the anticipated lack of significant population differentiation is a result of the wind-pollination characteristic of all known cannabis cultivars, coupled with their notable heterozygosity. To ensure the preservation of their genetic uniformity, numerous marijuana strains are propagated clonally rather than through seed.

GWAS has previously been successful in identifying genotype–phenotype associations in hop (*Humulus lupulus*) and cannabis (Henning et al., 2019). Moreover, one of the earliest studies of the cannabis genome and transcriptome found that the genes responsible for producing THC and CBD were located in different regions of the genome than previously thought and that there have been extensive rearrangements and variations in these regions among different cultivars of the plant (van Bakel et al., 2011). For marker–trait association in our study, multiple proteins across the genome are involved in traits related to cannabinoid yields, such as THC and CBD content. This implies that the production of cannabinoids is not only directly tied to the genes responsible for their synthesis in the cannabinoid pathway (Sawler et al., 2015; Hurgobin et al., 2021), but rather that additional areas of the genome also control these biosynthetic pathways (Marks et al., 2009; Laverty et al., 2019; Welling et al., 2020). This finding aligns with the earlier studies, further supporting the notion that multiple genetic factors contribute to cannabinoid content variation (Stout et al., 2012; Weiblen et al., 2015; Grassa et al., 2018; Braich et al., 2019; Zager et al., 2019; Livingston et al., 2020). Here we contribute further to this literature by confirming some of these important loci, but also identifying other novel loci for targeted future selection and breeding.

Furthermore, an additional SNP marker -816770_13-located on chromosome 1 and associated with the locus LOC115705717 (RefSeq accession: XM_030633131.1), responsible for 3-ketoacyl-CoA synthesis 19, has been found to be linked with CBD concentration. This SNP was previously identified as being connected to Autoflower1, which is involved in regulating the flowering time of hemp (Toth et al., 2022). These findings suggest that certain genes involved in the regulation of flowering time may also be correlated with cannabinoid content. One limitation of this study is that the phenotypic data collected here are only reported for a single growing season and environment, and thus, some caution must be used in generalizing the importance of these findings. However, given that our study confirms earlier identified genetic loci, such as those for flowering linked to cannabinoid production, this gives us confidence that our approach, is worthy and valid but should, in the future, be complemented with additional field trials and analyses to confirm these and other genetic loci.

Regarding the sex trait, the significant SNPs were annotated to uncover potential genetic mechanisms associated with sex determination. While a total of eight potential SNPs located on different chromosomes were identified for this trait, the prior study by Soorni *et al*. (2017), which failed to pinpoint distinct alleles for the regions responsible for sex determination. As the results of this study and earlier studies showed, the process of sex determination in cannabis is intricate and is not solely related to sex chromosomes (McKernan et al., 2020). It appears that the sex determination mechanism may be influenced by environmental factors and chemical applications (Lubell and Brand, 2018; Campbell et al., 2021) and involves the participation of other candidate genes like those related to trichome growth, sex determination, hermaphroditism, and photoperiod independence (Hurgobin et al., 2021) or genes involved in regulating phytohormone balance and the development of male flowers in female plants (Petit et al., 2020a). For example, the gene LOC115720754 (RefSeq accession: XM_030649931.1), located on chromosome 2, encodes the Zinc finger protein GAI-ASSOCIATED FACTOR 1, which acts as a transcription factor and a positive regulator of gibberellin (GA) action, homeostasis, and signaling in Arabidopsis (Fukazawa et al., 2014). This candidate gene was found to be associated with flowering time (Colasanti et al., 2006). In plants, GA has a role in flowering control (Goldberg-Moeller et al., 2013), and Petit *et al*. (2020b) linked the control of the flowering pathways in cannabis to that of sex determination, further highlighting the complexity of these inter-linked traits, where further research is warranted.

Out of the eight candidate loci identified for plant height, we observed that LOC115709353 (RefSeq accession: XM_030637438.1), located on chromosome 3, encodes anaphase-promoting complex subunit 8. In Arabidopsis, this protein is known to play a role in various aspects of development and embryogenesis by regulating the cell cycle, cell division, cell elongation, and endoreduplication control (Eloy et al., 2012; Xu et al., 2019; Saleme et al., 2021). While QTL analysis has been previously conducted for a range of agronomic traits on a population of 375 individuals (Woods et al., 2021), it has not been performed for the unique array of morpho-physiological traits presented here. These triats include, the number of nodes, internode length, crown length, and the number of leaves, as well as inflorescence-related features such as the number of lateral inflorescences and inflorescences length. The identified SNP markers for these traits have not been previously mapped, and consequently, this study stands as the inaugural attempt to assess these specific traits through association analysis, with loci and candidate genes reported, using the recently available genome sequence of *C. sativa*. The significant SNPs identified are novel for a range of traits and are not shared among them. These SNPs have the potential to serve as markers for marker-assisted breeding in cannabis, pending proper validation.

## CONCLUSION

Using GBS data from a diverse Iranian cannabis collection of wild germplasm (CGRC), this study has provided significant insights into the genetic variation and differentiation, population structure, and genotype-phenotype associations for this novel germplasm and how it differs from currently available global hemp and marijuana collections. Population structure analysis revealed five distinct groups in the Iranian cannabis collection. Pairwise F_ST_ comparisons identified the northeast population of Iran as the most genetically distinct, making it a priority for future breeding programs. Furthermore, the study confirmed several gene targets for unique traits, including inflorescence features, flowering time, cannabinoid content, sex, and some morphological traits. Together, this study has created a research platform that can link genomic variation and germplasm collection, facilitating selections for molecular breeding. These findings have important implications for improving the quality and productivity of new commercial cannabis varieties through breeding.

## FUNDING

This research was funded by BRC (Biopharmaceutical Research Company) as part of a grant to GT at UC Davis and by the Iran National Science Foundation (INSF) under grant number 96014753. Also, the research in the laboratory of GT is supported by the John B Orr endowment.

## ACKNOWLEDGMENTS

The authors thank the Iran National Science Foundation (INSF) for their support. The authors also express their profound gratitude to Afshin Peirovi, CIAN Diagnostics, 5330 Spectrum Drive, Suite I, Frederick, MD 21703, USA, for invaluable assistance in the preparation of lyophilized samples.

## Conflict of Interest

Author Annabelle Damerum is employed by Zymo Research Corporation, and authors George Hodgin and Brian Brandley are employed by Biopharmaceutical Research Company. The remaining authors declare that the research was conducted in the absence of any commercial or financial relationships that could be construed as a potential conflict of interest.

---

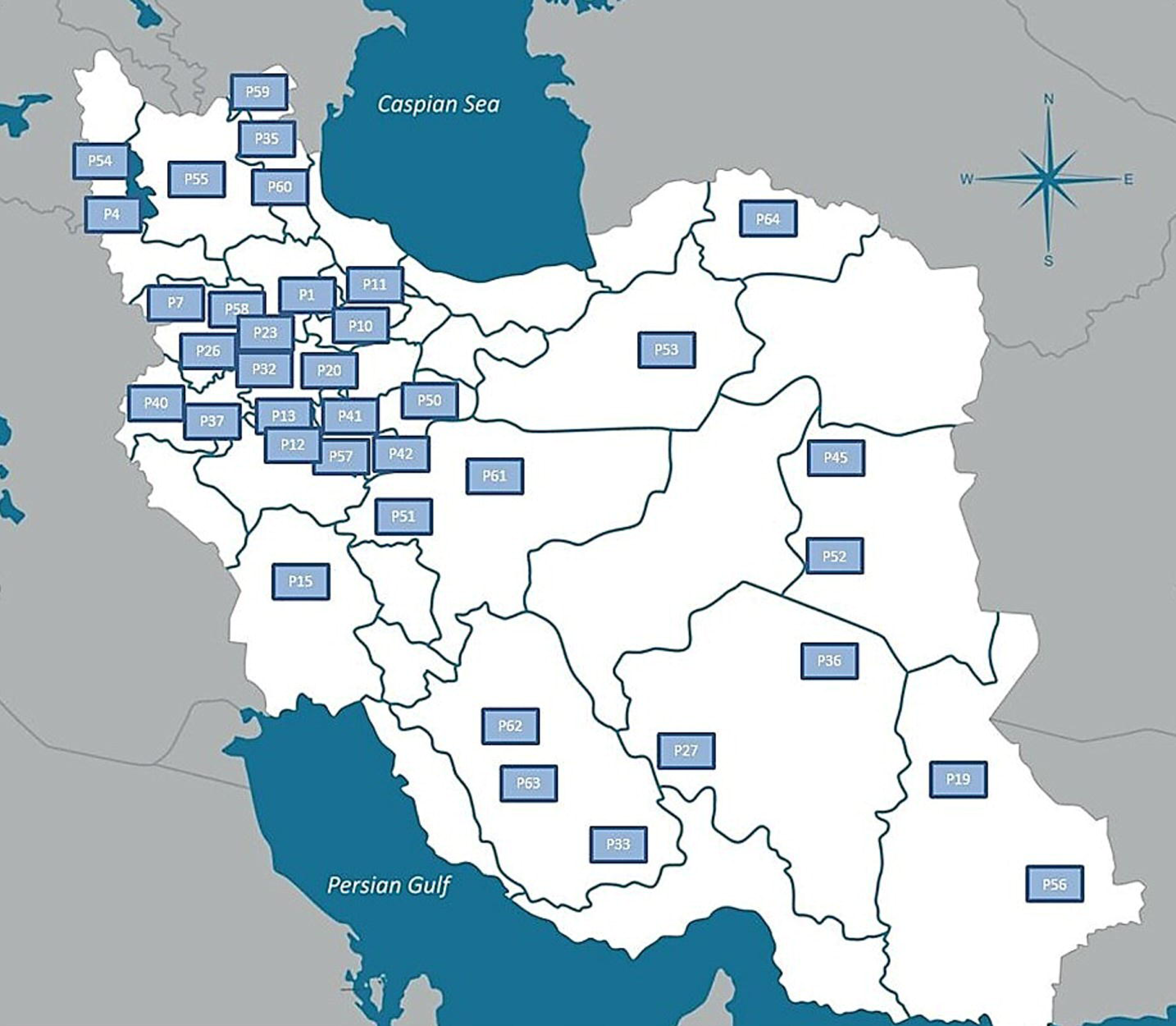

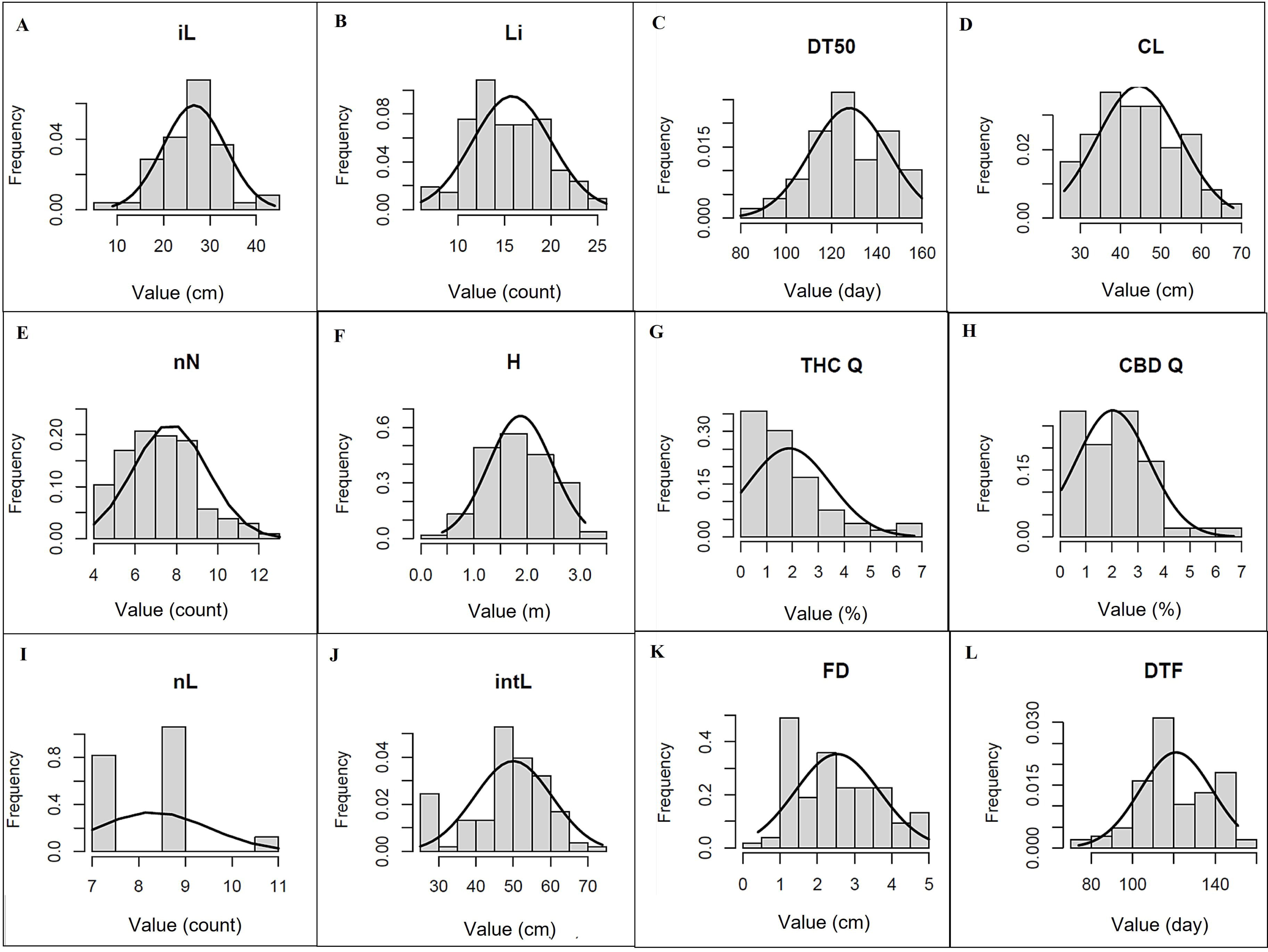

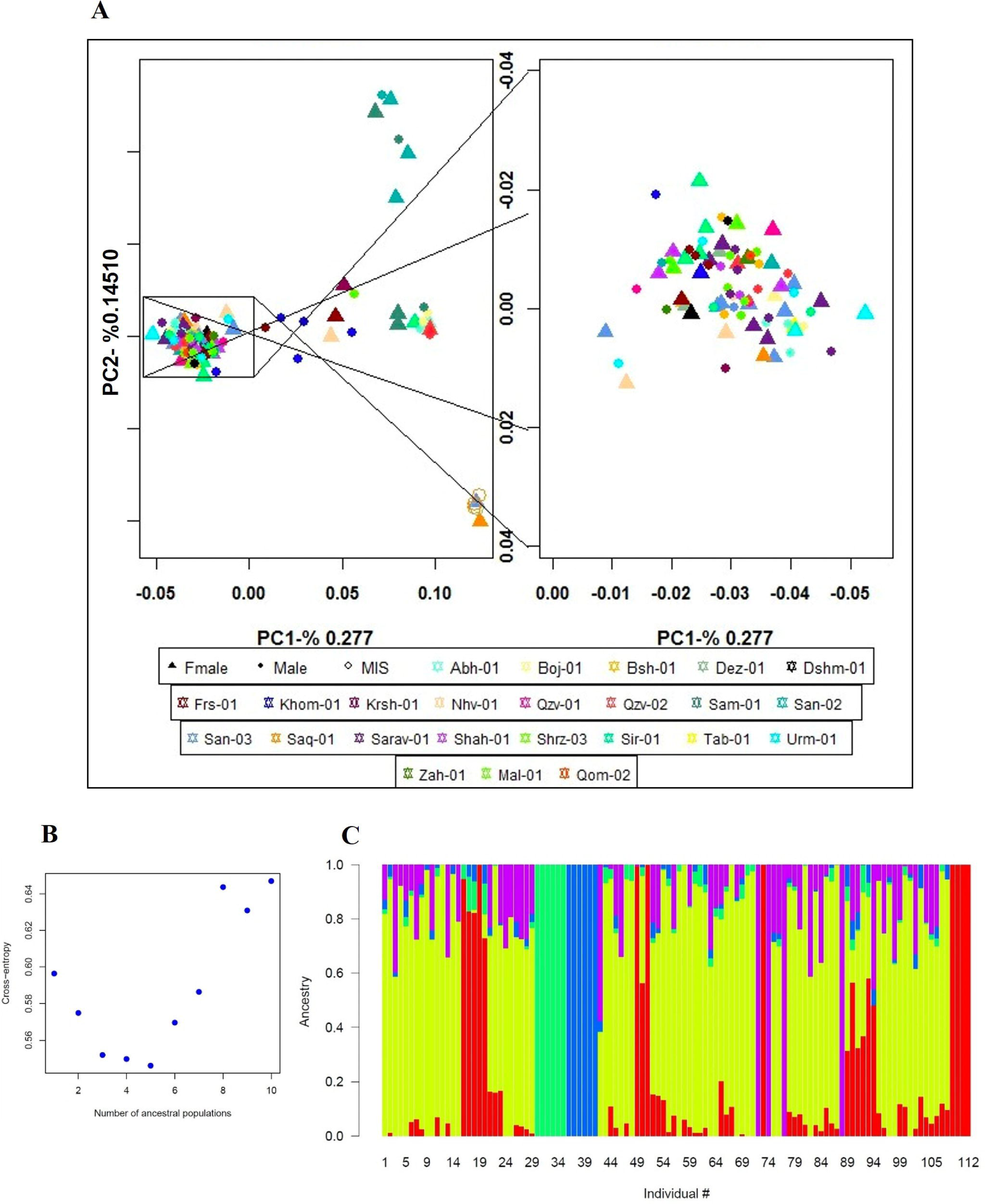

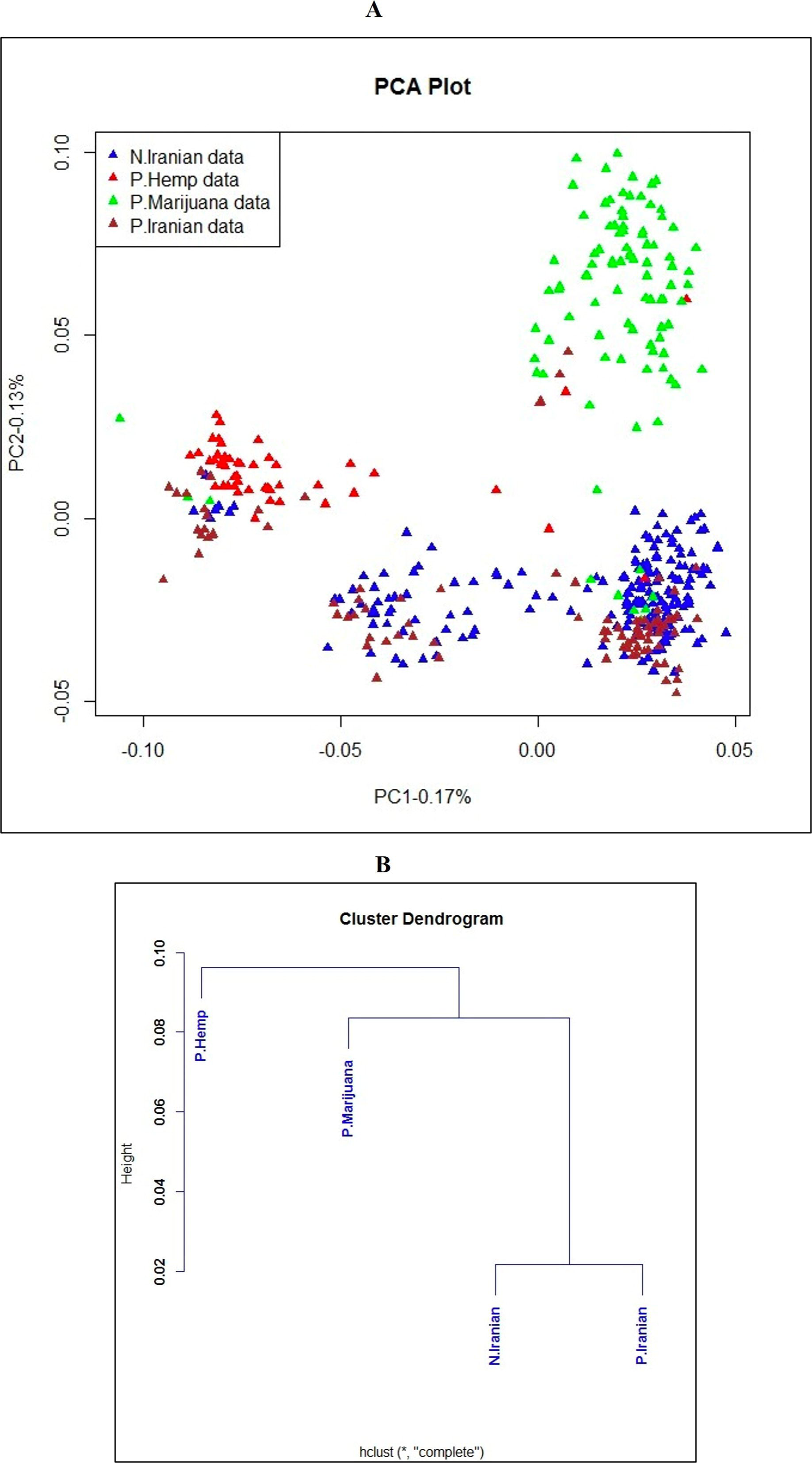

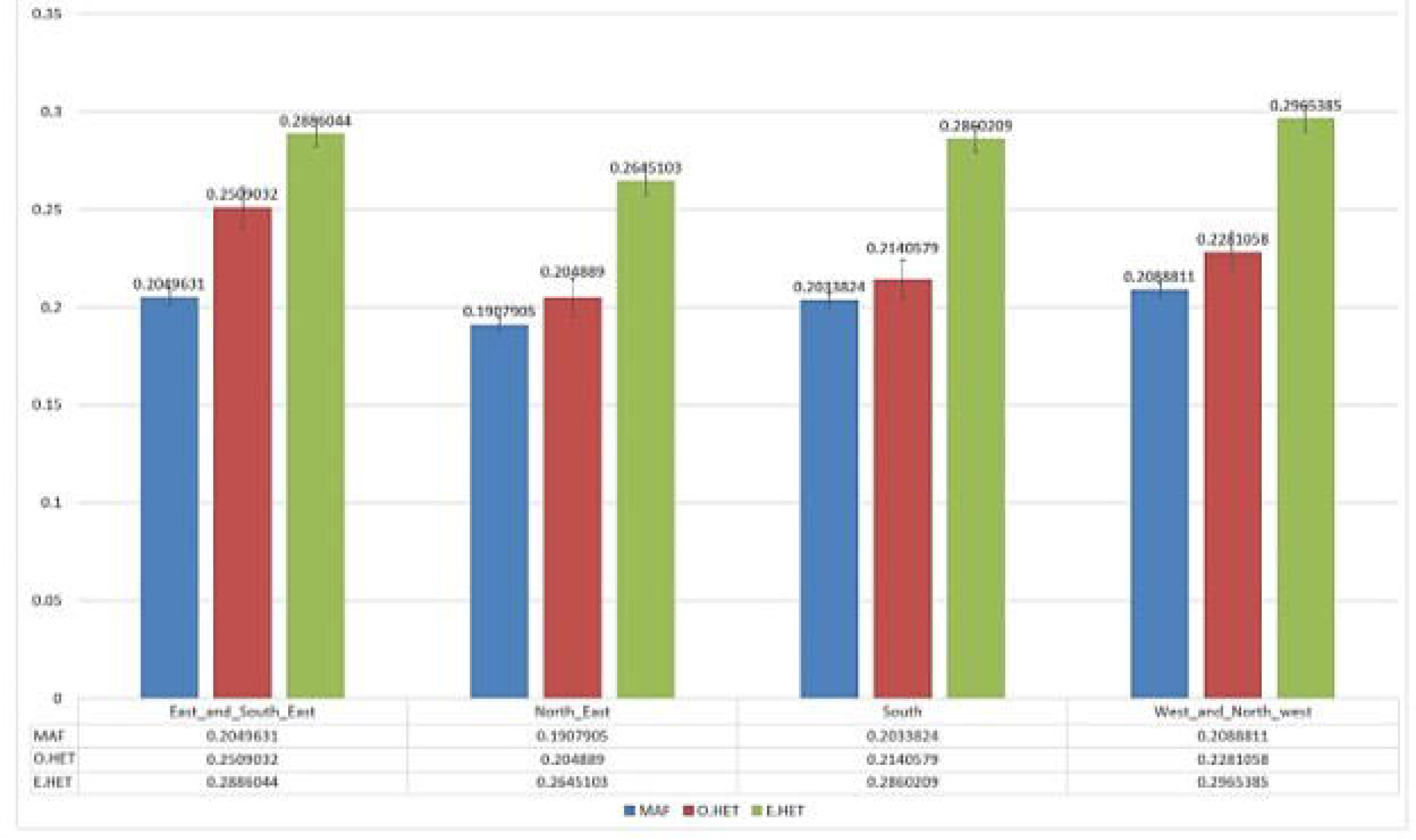

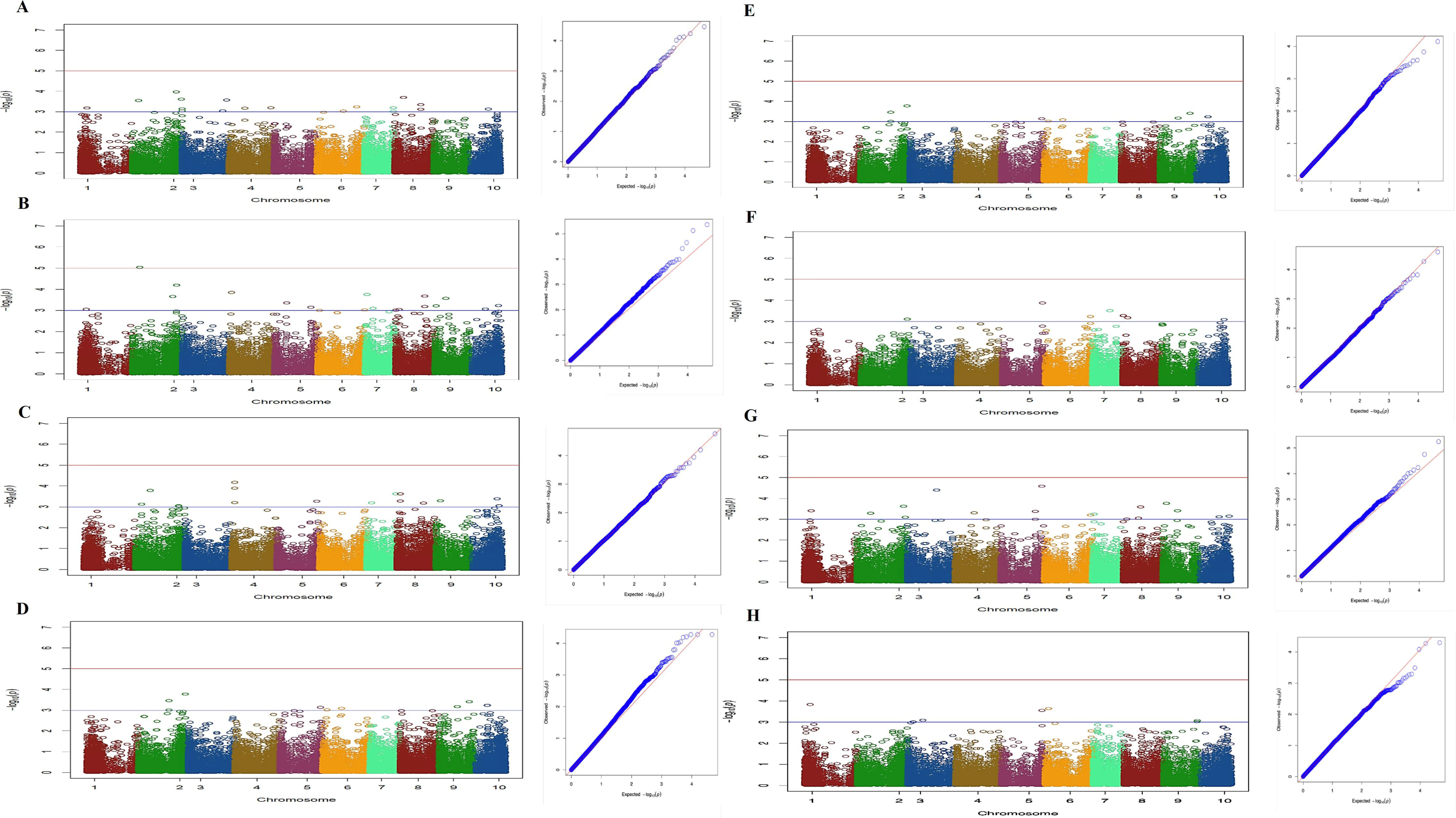

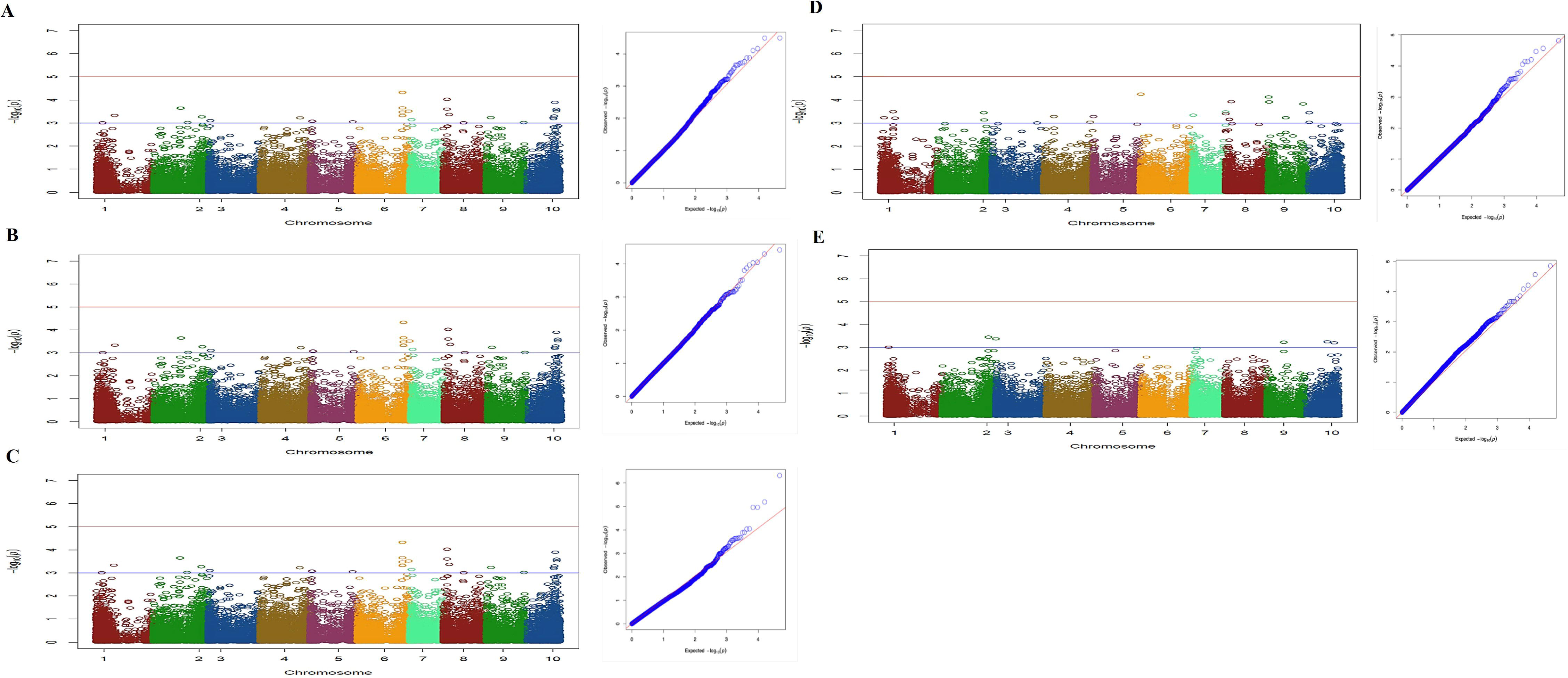

## Notes

### Competing Interest Statement

The authors have declared no competing interest.

