## Supplemental Data 1 for "Population genomics of a natural *Cannabis sativa* L. collection from Iran identifies novel genetic loci for flowering time, morphology, sex and chemotyping": Supplementary Figures.docx

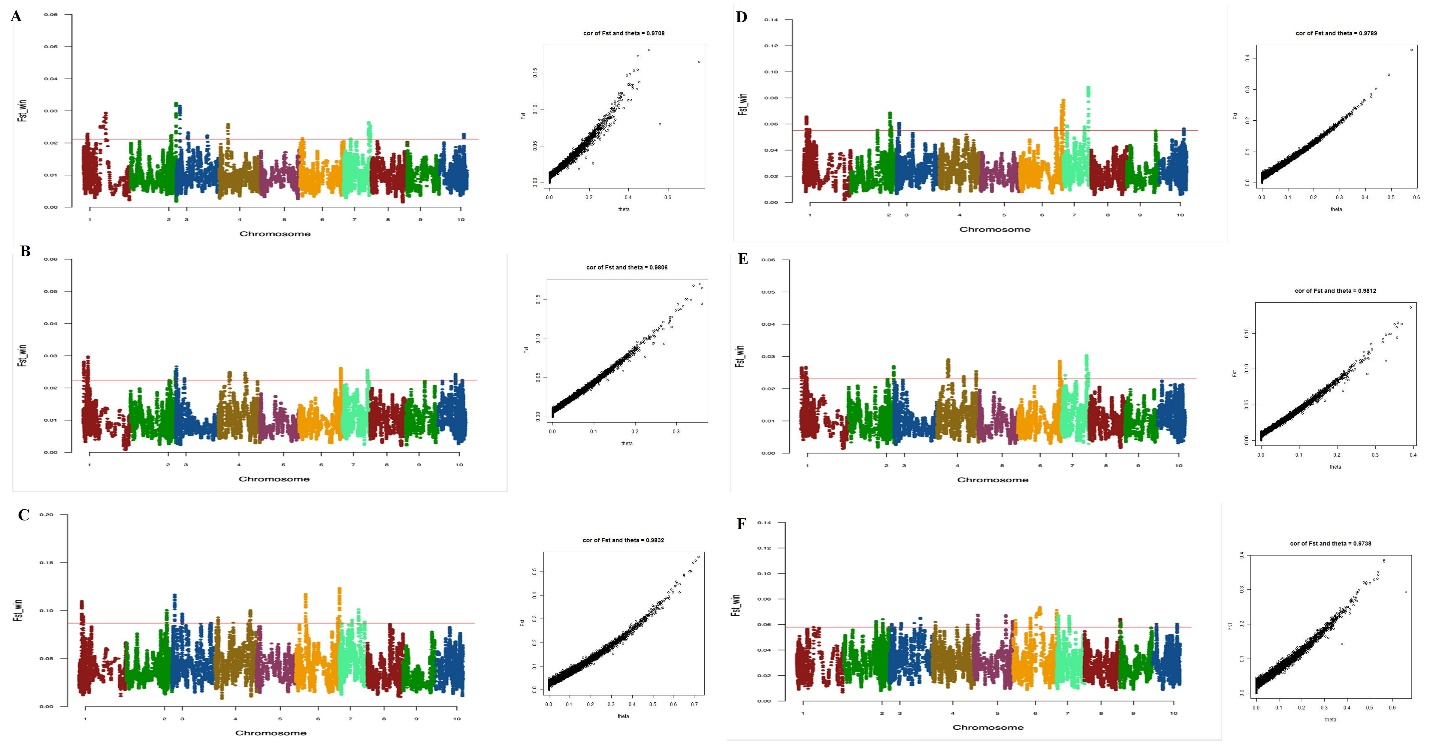

**Supplementary Figure 1.** Pairwise F_ST_ Manhattan plots for studied Iranian populations classified based on geographical distribution and climatic variables: (**A**) northeast: west and northwest (**B**) south: west and northwest (**C**) east and southeast: northeast (**D**) east and southeast: south (**E**) east and southeast: west and northwest (**F**) northeast: south.

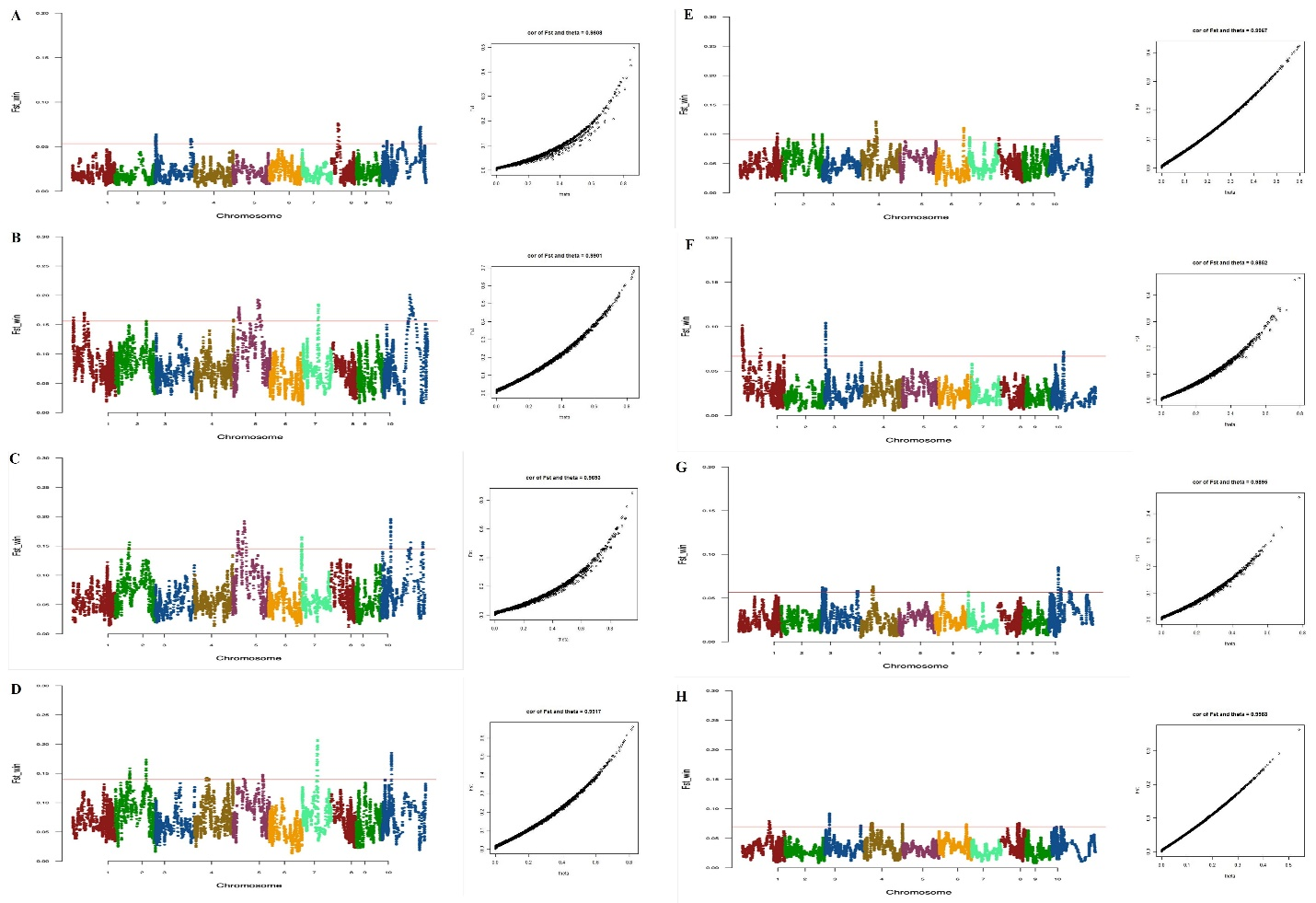
**Supplementary Figure 2.** Pairwise F_ST_ Manhattan plots for four geography-based Iranian populations and previously studied Iranian samples and global collections including hemp and marijuana. (**A**) marijuana population: northeast (**B**) hemp population: east and southeast (**C**) hemp population: northeast (**D**) hemp population: south (**E**) hemp population: west and northwest (**F**) marijuana population: east and southeast (**G**) marijuana population: south (**H**) marijuana population: west and northwest.

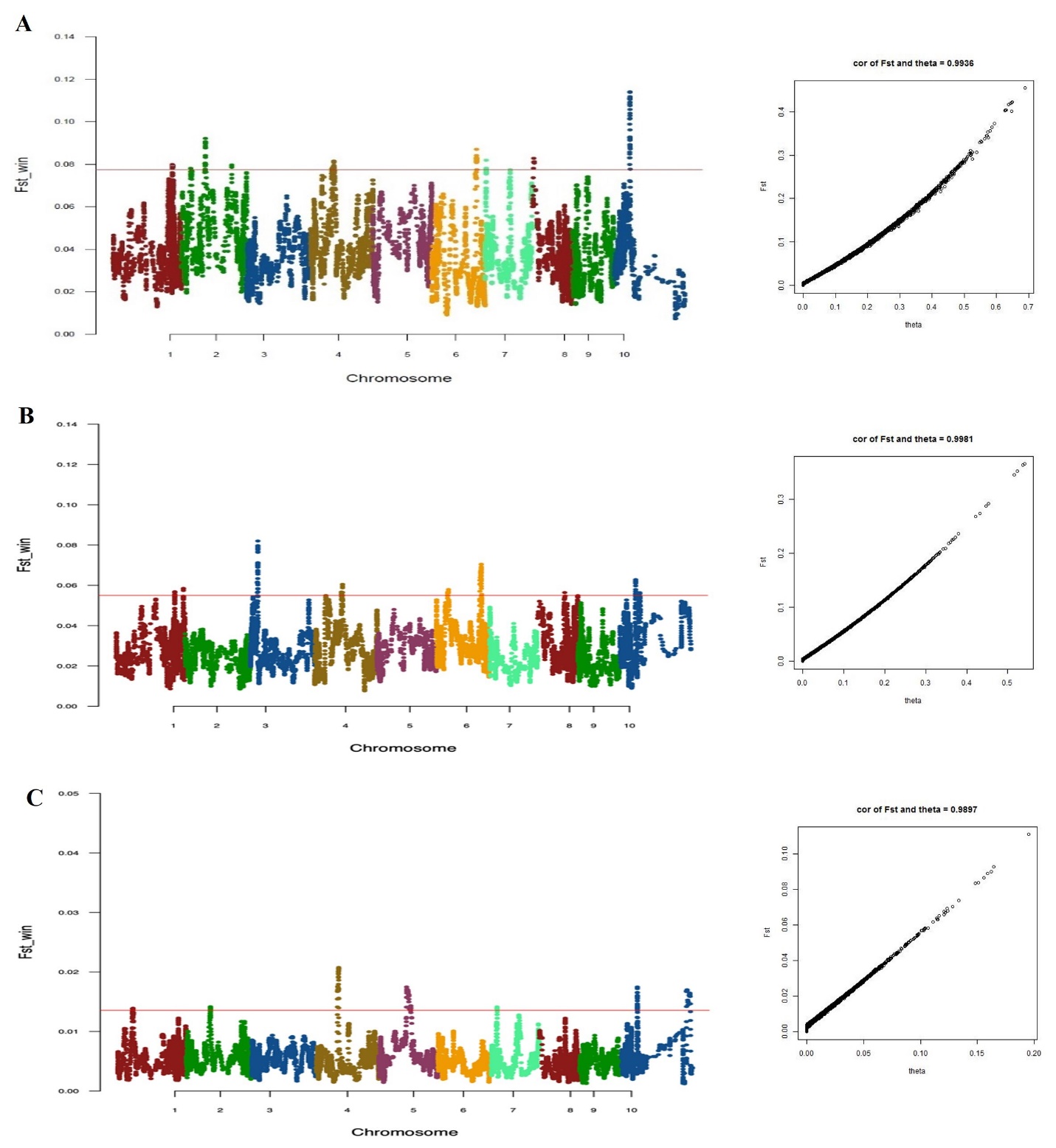

**Supplementary Figure 3.** Pairwise F_ST_ Manhattan plots for Iranian samples of this study as a single population and hemp/marijuana and previously studied Iranian samples. (**A**) Iranian samples of current study: hemp population (**B**) Iranian samples of current study: marijuana population (**C**) Iranian samples of current study: previously studied Iranian samples.
